# Novel strains of *Klebsiella africana* and *Klebsiella pneumoniae* in Australian Fruit Bats (*Pteropus poliocephalus*)

**DOI:** 10.1101/2021.07.12.451971

**Authors:** Fiona K. McDougall, Kelly L. Wyres, Louise M. Judd, Wayne S.J. Boardman, Kathryn E. Holt, Michelle L. Power

## Abstract

Over the past decade human associated multidrug resistant (MDR) and hypervirulent *Klebsiella pneumoniae* lineages have been increasingly detected in wildlife. This study investigated the occurrence of *K. pneumoniae* species complex (KpSC) in grey-headed flying foxes (GHFF), an Australian fruit bat. Thirty-nine KpSC isolates were cultured from 275 GHFF faecal samples (14.2%), comprising *K. pneumoniae* (*sensu stricto*) (*n*=30), *Klebsiella africana* (*n*=8) and *Klebsiella variicola* subsp. *variicola* (*n*=1). The majority (79.5%) of isolates belonged to novel sequence types (ST), including two novel *K. africana* STs. This is the first report of *K. africana* outside of Africa and in a non-human host. A minority (15.4%) of GHFF KpSC isolates shared STs with human clinical *K. pneumoniae* strains, of which, none belonged to MDR clonal lineages that cause frequent nosocomial outbreaks, and no isolates were characterised as hypervirulent. The occurrence of KpSC isolates carrying acquired antimicrobial resistance genes in GHFF was low (1.1%), with three *K. pneumoniae* isolates harbouring both fluoroquinolone and trimethoprim resistance genes. This study indicates that GHFF are not reservoirs for MDR and hypervirulent KpSC strains, but they do carry novel *K. africana* lineages. The health risks associated with KpSC carriage by GHFF are deemed low for the public and GHFF.

## 1. Introduction

*Klebsiella pneumoniae* is both a commensal and an opportunistic pathogen of humans and other vertebrate hosts, and a leading cause of human nosocomial and community acquired infections [1, 2]. The emergence and spread of multidrug resistant (MDR) *K. pneumoniae* strains exhibiting resistance to critically important antimicrobials [3, 4], specifically extended-spectrum beta-lactamase (ESBL) and carbapenemase producers, are considered an important global public health threat [5, 6]. Increasing reports of wildlife harbouring *K. pneumoniae* exhibiting multidrug resistance, including to carbapenems, extended-spectrum beta-lactams and fluoroquinolones [7-10] may have important implications for the spread of MDR *K. pneumoniae*. Birds in particular, have been identified as reservoirs and potential long-distance disseminators of ESBL producing *K. pneumoniae* [11-14]. Bats, like birds, are capable of flying long-distances [15, 16] and have the potential to be reservoirs and disseminators of bacterial pathogens, including *K. pneumoniae* [17-21] and any associated antimicrobial resistance (AMR) traits [22].

Whole genome sequencing (WGS) has revealed that isolates identified as *K. pneumoniae* through standard biochemical or mass spectrometry techniques actually comprise seven distinct taxa, informally referred to as the *K. pneumoniae* species complex (KpSC) [23]. The seven taxa forming KpSC are *K. pneumoniae* (*sensu stricto*) (Kp1), *K. quasipneumoniae* subsp. *quasipneumoniae* (Kp2), *K. quasipneumoniae* subsp. *similipneumoniae* (Kp4), *K. variicola* subsp. *variicola* (Kp3), and the more recently described *K. variicola* subsp. *tropica* (Kp5), *K. quasivariicola* (Kp6) and *K. africana* (Kp7) [24]. KpSC strains can be differentiated and assigned to unique sequence types (ST) using multilocus sequence typing (MLST) of seven housekeeping genes [25, 26].

The majority (∼85%) of KpSC isolates associated with clinical disease belong to *K. pneumoniae* (*sensu stricto*) [23], however, *K. variicola* subsp. *variicola* and *K. quasipneumoniae* are classified as emerging human pathogens [27, 28]. In contrast, the recently described *K. quasivariicola* has been reported in only a few clinical cases, and *K. africana* in a single human clinical case [24]. *Klebsiella pneumoniae* hospital outbreaks are predominantly caused by MDR clonal lineages, including clonal groups (CG) comprised of closely related STs, with CG258 (ST11, ST258, ST340, ST437 and ST512) and CG15 (ST14 and ST15) among the most common [29]. In recent decades, ‘hypervirulent’ *K. pneumoniae* have undergone global dissemination, becoming a leading cause of severe invasive community acquired infections [2]. The majority of hypervirulent *K. pneumoniae* belong to a handful of clones, the most common being ST23, ST65 and ST86 [30]. The hypervirulent phenotype is associated with the presence of specific pathogenicity and virulence factors, including capsule types K1 and K2, serotypes O1 and O2, capsule upregulator genes *rmpA* and *rmpA2*, and siderophores yersiniabactin, aerobactin, salmochelin and colibactin [30]. Whilst MDR and hypervirulent *K. pneumoniae* are usually distinct strains, there are recent reports of convergence, with isolates exhibiting both MDR and hypervirulence traits [23]. *Klebsiella* is intrinsically resistant to ampicillin and amoxicillin due to the presence of core chromosomal beta-lactamase (*bla*) producing genes [6], designated *bla*_SHV_ in *K. pneumoniae, bla*_LEN_ in *K. variicola* and *K. quasivariicola*, and *bla*_OKP_ in other KpSC [6, 24, 31, 32]. Acquired AMR in *K. pneumoniae* is typically associated with acquisition of resistance genes via horizontal transfer of mobile genetic elements, such as plasmids and transposons [33] and integrons [34]. One type of integron, the clinical class 1 integron, has been significant for the emergence and on-going dissemination of AMR in Gram-negative bacteria including *K. pneumoniae* [35]. Class 1 integrons capture and express diverse antibiotic resistance genes (ARGs) via the integrase gene (*intl1*) and a promoter (*Pc*), and are typically characterised by a 3’-conserved segment (*qacEΔ1-sul1*) [34]. *K. pneumoniae* is proposed to play an important role in the acquisition of AMR genes from environmental bacteria and horizontal transmission via mobile genetic elements to other Enterobacteriaceae [35, 36]. Notably, the emergence and dissemination of plasmid-borne *bla*_SHV_ genes associated with ESBL activity [37, 38] and carbapenemase genes (*bla*_NDM-1_, *bla*_KPC_, *bla*_OXA-48_) [36], are now widely found in diverse Enterobacteriaceae.

The One Health approach to AMR highlights the need for ecological studies of *K. pneumoniae* in diverse animal species to identify reservoirs of clinically important lineages [23]. Surveillance of wildlife for pathogenic and antimicrobial resistant bacteria is particularly important for species inhabiting urban environments, such as bats, due to the zoonotic risks posed to humans in close proximity.

Bats belong to the large speciose order Chiroptera and are widely distributed around the world with many species exploiting urban environments [39], and they are known reservoirs of ESBL and carbapenemase producing *K. pneumoniae*. For example, ESBL producing *K. pneumoniae* (*bla*_CTX-M-15_) were detected in Franquet’s epauletted fruit bats (*Epomops franqueti*) and Woermann’s fruit bats (*Megaloglossus woermanni*) in Gabon [17] and carbapenemase producing *K. pneumoniae* harbouring *bla*_OXA-48_ (ST1878) and *bla*_KPC-3_ (ST512) in microbat guano from Algeria [18].

Class 1 integrons harbouring ARGs to narrow-spectrum penicillins, trimethoprim and aminoglycosides were detected in faecal DNA from grey-headed flying fox (GHFF; *Pteropus poliocephalus*) a large fruit bat endemic to Australia [22], however, the bacterial hosts of these integrons have not been resolved. This study aimed to examine *K. pneumoniae* ecology in GHFF and to define the diversity, pathogenicity and virulence factor occurrence, and AMR gene carriage in isolates from GHFF.

## 2. Materials and methods

### 2.1. Sample collection

GHFF have established large colonies, comprising several thousand to >50,000 individuals, in both urban and rural environments [40, 41]. The distribution of GHFF spans eastern Australia, from Adelaide (South Australia) to Ingham (Queensland). Faecal samples (*n*=275) were collected between 2017 and 2018 from free-living wild GHFF (*n*=255) inhabiting urban environments and captive GHFF (*n*=20) that were undergoing rehabilitation. GHFF faecal samples were collected from four wild colonies; Camellia Gardens (CG), (*n*=50), Centennial Park (CP) (*n*=52) and Blackalls Park (BP) (*n*=101) in New South Wales (NSW), and Adelaide Botanic Park (ABP) (*n*=52) in South Australia (SA). Sampled captive animals were from the ABP colony that were recovering in a rehabilitation facility after heat stress (Fauna Rescue of South Australia), Mylor, SA (*n*=20). Faecal samples were collected either opportunistically from plastic drop sheets placed under roosting flying foxes or directly from individual GHFF via a rectal swab for the ABP colony. The FecalSwab™ (COPAN, Brescia, Italy) system was used for both sampling methods. Samples were stored at 4°C and cultured within 72 h of collection.

### 2.2. Ethics

All sample collections were conducted under approvals from animal ethics committees at Macquarie University (No. 2017/013) and The University of Adelaide (No. S-2015-028), NSW Government Scientific Licence (No. SL101898) and SA Department of Environment and Water Wildlife Scientific Permit (M-23671-1,2 and 3).

### 2.3. Klebsiella isolation and DNA extraction

FecalSwab media (200μL) was inoculated into 5 mL of Luria-Bertani (LB) broth (Difco Laboratories, Detroit, USA) supplemented with amoxicillin (10 mg/L) (Sigma-Aldrich, St. Louis, USA) and incubated overnight at 37°C. Broth cultures were then streaked onto modified SCAI medium [42] which comprised of Simmons citrate agar (Oxoid, Hamphire, United Kingdom) supplemented with 1% myo-inositol (Sigma, St. Louis, USA) and amoxicillin (10 mg/L) and incubated at 37°C for 40-48 h. Yellow moist colonies indicating *Klebsiella* sp., *Enterobacter* sp. or *Serratia* sp. were selected for further analyses. To distinguish *Klebsiella* sp. from *Serratia* sp., isolates were evaluated for gelatin metabolism by inoculation into 5 mL of nutrient gelatin (NG) containing 13 g/L nutrient broth (Oxoid, Hamphire, United Kingdom) and 120 g/L commercial bovine gelatine (Dr. Oetker, Bielefeld, Germany). The NG samples were incubated for 48-96 h and tested after 48 h, 72 h and 96 h for a positive gelatin metabolism reaction by transferring to 4°C for 30 min. Isolates that were negative for gelatin metabolism (solid at 4°C) after 96 h were deemed presumptive *Klebsiella* sp. or *Enterobacter* sp. with those positive for gelatin metabolism (liquid at 4°C) eliminated from further analyses. Bacterial cell lysate (95°C 15 min; >10000 g 10 min) was used as the DNA template in KpSC MLST and class 1 integron PCRs.

### 2.4. Preliminary identification of KpSC

Preliminary identification of KpSC isolates was performed by MLST PCRs targeting three of seven house-keeping genes (*gapA, mdh* and *phoE*) using primer sets listed in Table S1 [25]. DNA from *K. pneumoniae* strain DAR_Y9835_6192 was used as a positive control in all PCRs. PCRs were performed using GoTaq® Colorless Mastermix (Promega, Madison, USA), 0.4 μM of each primer and 2 μL DNA. The PCR conditions were 94°C for 2 min, then 35 cycles of 94°C for 20 s and a primer specific annealing temperature for 30 s (Table S1) and extension at 72°C for 30 s, with a final extension at 72°C for 5 min [43]. Isolates returning a positive result for all three housekeeping genes when analysed by agarose gel electrophoresis were presumed to be *K. pneumoniae* and underwent whole genome sequencing and class 1 integron screening.

### 2.5. Screening for class 1 integrons

Isolates were screened for the class 1 integron integrase gene (*intI1*) using primers HS463a and HS464 (Table S1) and *IntI1* positive isolates were then amplified using primers HS458 and HS459 (target the conserved *attl1* and 3’ *qacEΔ1* region of the class 1 integron) (Table S1). All integron PCRs were performed using GoTaq® Colorless Mastermix (Promega, Madison, Wisconsin, USA), 0.4 μM of each primer and 2 μL DNA (boil preparation method) as previously described [22].

### 2.6. Whole genome sequencing and genetic characterisation

DNA was extracted using the GenFind v2 kit (Beckman Coulter) and libraries prepared using the NexteraFlex kit (Illumina Inc.) as per the manufacturers’ guidelines. Libraries were sequenced on the Illumina NextSeq500 platform (150bp paired-end reads, n=52 presumptive KpSC). Following preliminary analysis, a single *K. africana* isolate (FF1003) was selected for additional long read sequencing using the Oxford Nanopore MinION R9 device (Oxford Nanopore Technologies) as described previously [44]. Illumina reads (of 150 bp) were quality trimmed using Cutadapt v1.16 [45] and *de novo* assembled using SPAdes v3.1.12 [46] optimised with Unicycler v0.4.7 [47]. The FF1003 genome was fully resolved using Unicycler’s hybrid assembly approach. Illumina reads were also mapped to the *K. pneumoniae* SGH10 reference chromosome (GenBank accession: CP025080) and single nucleotide variants called using the RedDog v1b 10.3 pipeline (https://github.com/katholt/RedDog). Genomes were excluded from further analysis if any of the following quality control criteria were met; i) total assembly size <5Mbp or >6Mbp; ii) mean read depth <20x (calculated from mapping stats); iii) mapping coverage <85%; iv) ratio of heterozygous to homozygous variant calls >0.4 (cut-off determined empirically); v) species not part of the KpSC (see below).

Genotypic characterisation, including species confirmation, KpSC MLST, as per the published scheme [25, 26], capsule (K) and lipopolysaccharide (O) locus types, ARGs and virulence genes, were determined from *de novo* assemblies using Kleborate v1.0 (https://github.com/katholt/Kleborate). Novel MLSTs were submitted for assignment at the *K. pneumoniae* BIGSdb hosted by the Institute Pasteur, France (https://bigsdb.pasteur.fr/klebsiella/). The genetic contexts of acquired AMR genes were investigated through manual inspection of the relevant assembly graphs using Bandage [48]. Plasmid replicon markers were identified using PlasmidFinder 2.1 [49] (available at https://cge.cbs.dtu.dk/services/PlasmidFinder/).

Maximum likelihood phylogenetic analysis was completed for all genomes passing quality control (n=39) using FastTree v2.1.20 [50] with an alignment of 404,516 core variant sites (present in ≥95% genomes, identified by RedDog using the SGH10 chromosomal reference; GenBank CP025080). An independent phylogenetic analysis comprising eight *K. africana* genomes from this study plus two published *K. africana* genomes [24, 51] and a *K. quasivariicola* outgroup (SRA accession; SRR5386043) was completed using RAxML v8.1.23 [52] (best of five runs each with 100 bootstrap replicates, general time reversible model with Gamma model of rate variation) with an alignment of 39,936 core variant sites (present in ≥95% genomes, identified by read-mapping to the FF1003 chromosome using RedDog). The phylogenetic trees were annotated using Interactive Tree of Life v5 (iTOL) (https://itol.embl.de) [53].

### 2.7. Data accessibility

All KpSC isolate WGS data is available in the NCBI Sequence Read Archive (SRA) under BioProject ID PRJNA646592 (BioSample accession numbers SAMN15547533 to SAMN15547571) and the assembled genome for *K. africana* isolate FF1003 is available in GenBank under accession numbers CP059391 to CP059393. Individual accession numbers are available in supplementary Tables S2 and S3.

### 2.8. Phenotypic antibiotic resistance testing

Isolates harbouring acquired AMR genes (*n*=3) (as identified by WGS) and representative isolates of all other KpSC STs (*n*=12) underwent phenotypic antibiotic susceptibility testing for ESBL activity and for resistance to quinolones/fluoroquinolones, trimethoprim +/-sulfamethoxazole and aminoglycosides (Table S4). Antibiotic susceptibility testing was performed according to the European Committee on Antimicrobial Susceptibility Testing (EUCAST) disk diffusion method [54] and isolates were evaluated as susceptible or resistant using EUCAST breakpoint criteria (v 9.0 available at http://www.eucast.org/clinical_breakpoints/) (Table S4). EUCAST breakpoints were unavailable for nalidixic acid and susceptibility was instead determined using the Clinical and Laboratory Standards Institute (CSLI) breakpoint criteria (CLSI M100 ED29:2019 available at https://clsi.org/standards/products/free-resources/access-our-free-resources/) (Table S4).

## 3. Results

### 3.1. Occurrence of KpSC in GHFF

Of 50 presumptive *K. pneumoniae* isolates identified by MLST, WGS confirmed 39 genomes passed quality control and belonged to the KpSC, with the majority belonging to *K. pneumoniae* (*n*=30) and the remaining to *K. africana* (*n*=8) and *K. variicola* subsp. *variicola* (*n*=1) (Fig. 1). The KpSC isolates were detected in three of four wild GHFF colonies, with frequencies ranging from 0.0% to 36.0% (overall mean 9.0%) (Fig. 1). For the captive GHFF, *K. pneumoniae* was detected in 80% of faecal samples from the single wildlife rehabilitation facility in this study (Fig. 1).

**Fig. 1.**
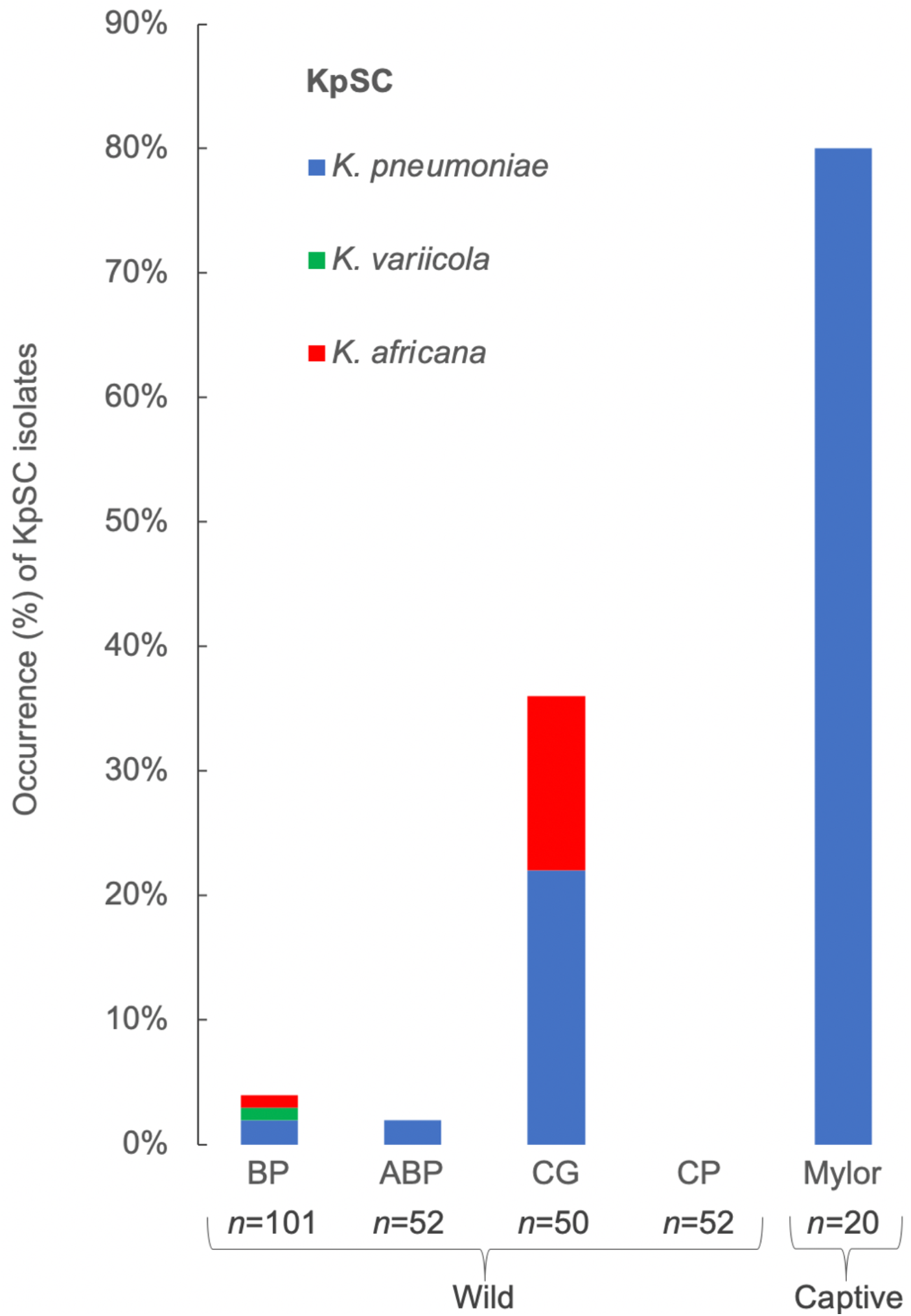
The occurrence and distribution of *K. pneumoniae* species complex isolates in GHFF from four wild colonies and one captive site. BP; Blackalls Park (NSW). CG; Camellia Gardens (Sydney, NSW). ABP; Adelaide Botanic Park (Adelaide, SA). CP; Centennial Park (Sydney, NSW). Mylor; Mylor captive GHFF rehabilitation facility (SA).

### 3.2. Phylogenetic diversity and geographical distribution

Diverse KpSC lineages, each associated with a distinct ST, were detected in GHFF across four sampling locations (Fig. 2). MLST identified 13 STs amongst the 39 GHFF KpSC isolates, which comprised eight novel STs (*K. pneumoniae, n*=5, *K. africana, n*=*2* and *K. variicola* subsp. *variicola, n=*1) and five recognised STs (ST105, ST661, ST1017, ST1412 and ST4919) (Fig. 2). The most frequent ST was the novel ST5035 (*n*=15 isolates). The eight *K. africana* isolates belonged to two novel STs that were designated ST4938 (*n*=7) and ST4939 (*n*=1) (Fig. 2). Eighteen isolates from the CG colony represented eight STs; *K. pneumoniae n*=6 STs and *K. africana n*=2 STs (Fig. 2). Four KpSC isolates were cultured from the BP colony, with two being distinct *K. pneumoniae* STs, one *K. africana* and one *K. variicola*, and one *K. pneumoniae* isolate was cultured from the ABP colony (Fig. 2). Although the captive Mylor GHFF showed a high occurrence of *K. pneumoniae*, 15 of 16 isolates were ST5035 and showed little genetic diversity (Fig. 2). Simpson diversity in the wild GHFF was 0.88 overall, which ranged from 0.84 in the CG colony to 1.0 in the BP colony, however, the captive GHFF group showed almost complete uniformity (Simpson diversity = 0.13).

**Fig. 2.**
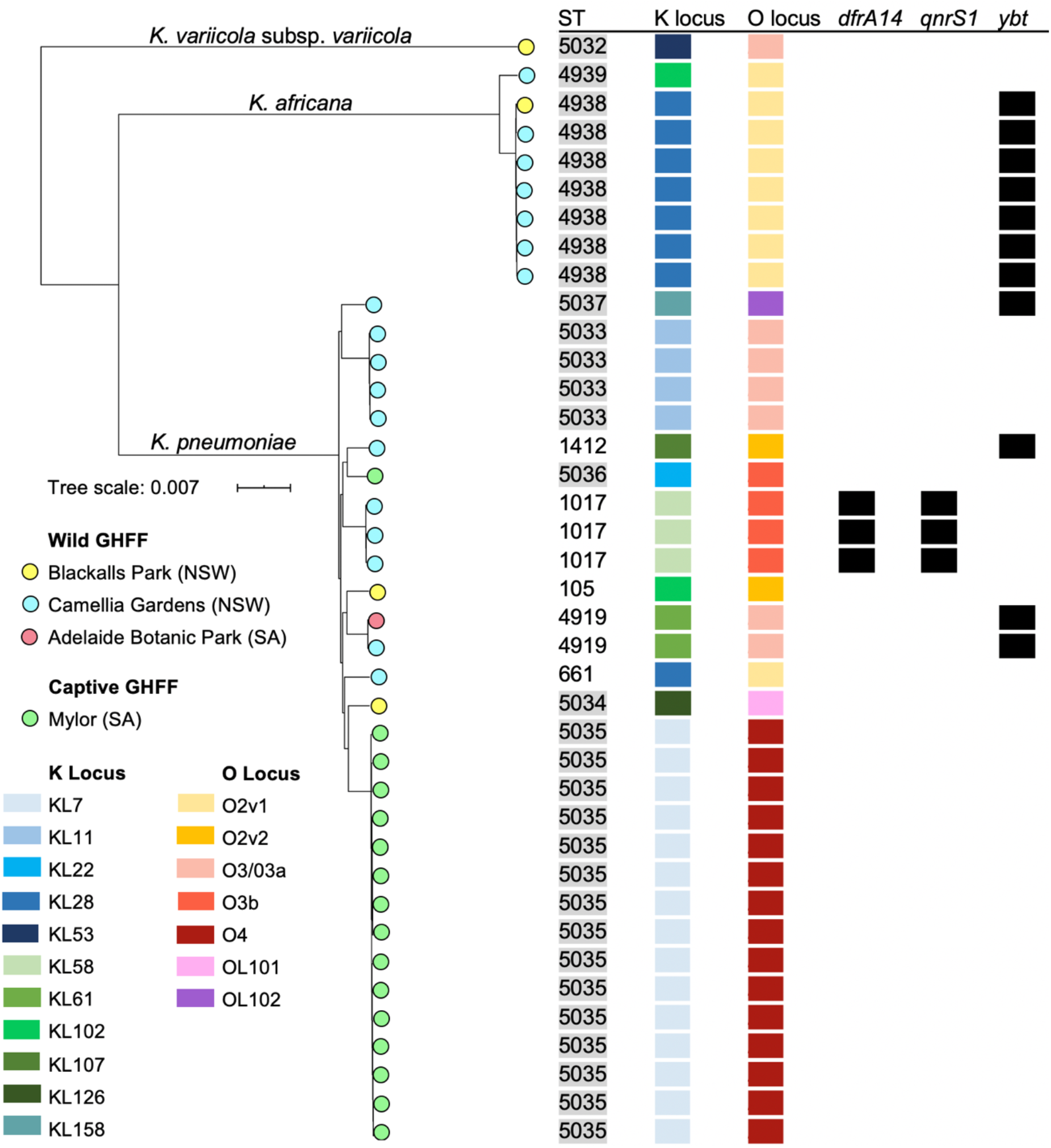
Core genome maximum likelihood phylogenetic tree constructed using FastTree showing the relatedness of all GHFF KpSC isolates with associated metadata. Single nucleotide variants (SNVs) were identified by RedDog using chromosomal reference SGH10 (GenBank CP025080). The tree is midpoint rooted. Scale bar indicates substitutions per site. Novel STs are highlighted in grey. Black squares indicate the presence of the antimicrobial resistance gene alleles *dfrA14* and *qnrS1*, and the yersiniabactin (*ybt*) virulence locus. ST, sequence-type.

STs were typically clustered by location, with the two notable exceptions being *K. africana* ST4938 which was detected in GHFF from the CG (*n*=6) and BP (*n*=1) colonies, and *K. pneumoniae* ST4919 in the ABP (*n*=1) and CG (*n*=1) colonies (Fig. 2).

### 3.3. K-loci and O-loci diversity

A total of 11 distinct K-loci were identified in the 39 GHFF KpSC isolates, with KL7 being the most frequently detected (*n*=15), occurring in all ST5035 isolates (Fig. 2). All *K. africana* ST4938 isolates (*n*=7) and ST661 (*n*=1) carried KL28, and the four ST5033 isolates carried KL11 (Fig. 2).

Five O-loci were identified, with O4 being the most frequent predicted serotype (*n*=15, all ST5035), followed by O2 (*n*=11, five different STs), O3 (*n*=11, five different STs), and two *K. pneumoniae* were associated with the less common OL101 and OL102 serotypes [55] (Fig. 2). Of the 11 KpSC isolates with the O2 serotype, the majority were associated with *K. africana* (*n*=8) and three were present in *K. pneumoniae* (ST105, ST661 and ST1412) (Fig. 2).

### 3.4. Intrinsic beta-lactam resistance genes

Amongst the *K. pneumoniae* isolates, the most frequent allele detected was a sequence variant of *bla*_SHV-110_ (as determined by the translated amino acid sequences) which was present in all ST5035 isolates (*n*=15). The remaining *K. pneumoniae* carried alleles with translated amino acid sequences identical to *bla*_SHV-1_ (*n*=8), *bla*_SHV-11.v1_ (*n*=3), *bla*_SHV-27_ (*n*=8) and one sequence variant of *bla*_SHV-110_. The single *K. variicola* isolate carried a *bla*_LEN-2_ variant and all *K. africana* isolates carried *bla*_OKP-C-1_ variants [24], as determined by the translated amino acid sequences. Phenotypic antimicrobial susceptibility testing confirmed all tested GHFF KpSC isolates (*n*=13) exhibited intrinsic resistance to amoxicillin and ampicillin, but not ESBL activity (Table S5).

### 3.5. Acquired antimicrobial resistance genes

Only one of 13 GHFF KpSC STs (ST1017) was associated with acquired ARGs, which equated to an overall occurrence of 1.1% across all sampled GHFF (wild 1.2% and captive 0.0%). ST1017 was cultured from three faecal samples from wild GHFF, with all three isolates carrying two AMR genes, *dfrA14* and *qnrS1*, which confer resistance to trimethoprim and fluoroquinolones respectively (Fig. 2). Phenotypic resistance to trimethoprim and ciprofloxacin, and intermediate resistance to nalidixic acid, was confirmed for all three ST1017 isolates by disk diffusion antibiotic susceptibility testing (Table S5).

Class 1 integron screening of GHFF KpSC isolates detected *IntI1* in all three ST1017 isolates however, amplification of the gene cassette array was not successful. BLASTn searches for these genes in the relevant genome assembly graphs indicated that q*nrS1, intI1* and *dfrA14* were co-harboured on a single replicon, which formed a closed circularised path through the graph which was subsequently shown to carry an IncN plasmid replicon marker, suggesting this structure represented an antimicrobial resistance plasmid. *DfrA14* was associated with *IntI1*, however, the class 1 integron was truncated and the 3’-conserved segment replaced by *mobC* and an IS*6100* transposase. A BLASTn search found the ST1017 *IntI1-drfA14-mobC-*IS*6100* integron sequence was an identical match to >100 *K. pneumoniae* associated plasmids in GenBank, including pathogenic strains (accessions CP052219, CP052525 and MN218814). The *qnrS1* gene was associated with an IS*26* transposase, with both inserted into a *tra* operon.

No KpSC STs (0/13) harboured acquired AMR genes conferring ESBL or carbapenemase activity, with all tested isolates susceptible to amoxicillin plus clavulanic acid, cephalexin, cefotaxime and imipenem in disk diffusion antibiotic susceptibility testing (Table S5).

### 3.6. Virulence factors

Kleborate searches resulted in the identification of one acquired siderophore, yersiniabactin, in 28.2% of GHFF KpSC isolates, including the seven ST4938 *K. africana* isolates (Fig. 2). Other virulence factors including aerobactin (*iuc*), salmochelin (*iro*), colibactin (*clb*), *rmpA, rmpA2* and capsule types K1 and K2 were not identified in any GHFF KpSC isolates.

### 3.7. Phylogenetic comparison of human and GHFF K. africana isolates

The GHFF *K. africana* isolates were compared to the only two previously reported *K. africana* isolates; a MDR clinical isolate (ERR315152, ST2831) causing a disseminated infection in Kenya and a human faecal isolate (ERR2835900, ST3291) from Senegal [24] (Fig. 3). The human isolates were separated by 20,512 chromosomal single nucleotide variants (SNVs) and both were distantly related to the GHFF isolates (≥16,249 pairwise chromosomal SNVs). However, seven of the eight GHFF isolates were clustered together with little genetic diversity evident between them (≤53 SNVs) (Fig. 3). Interestingly, these included isolates from two different sites (Camelia Gardens, *n*=6 and Blackhall’s Park, *n*=1). The eight GHFF isolates differed from the rest by ∼23,300 SNVs.

**Fig. 3.**
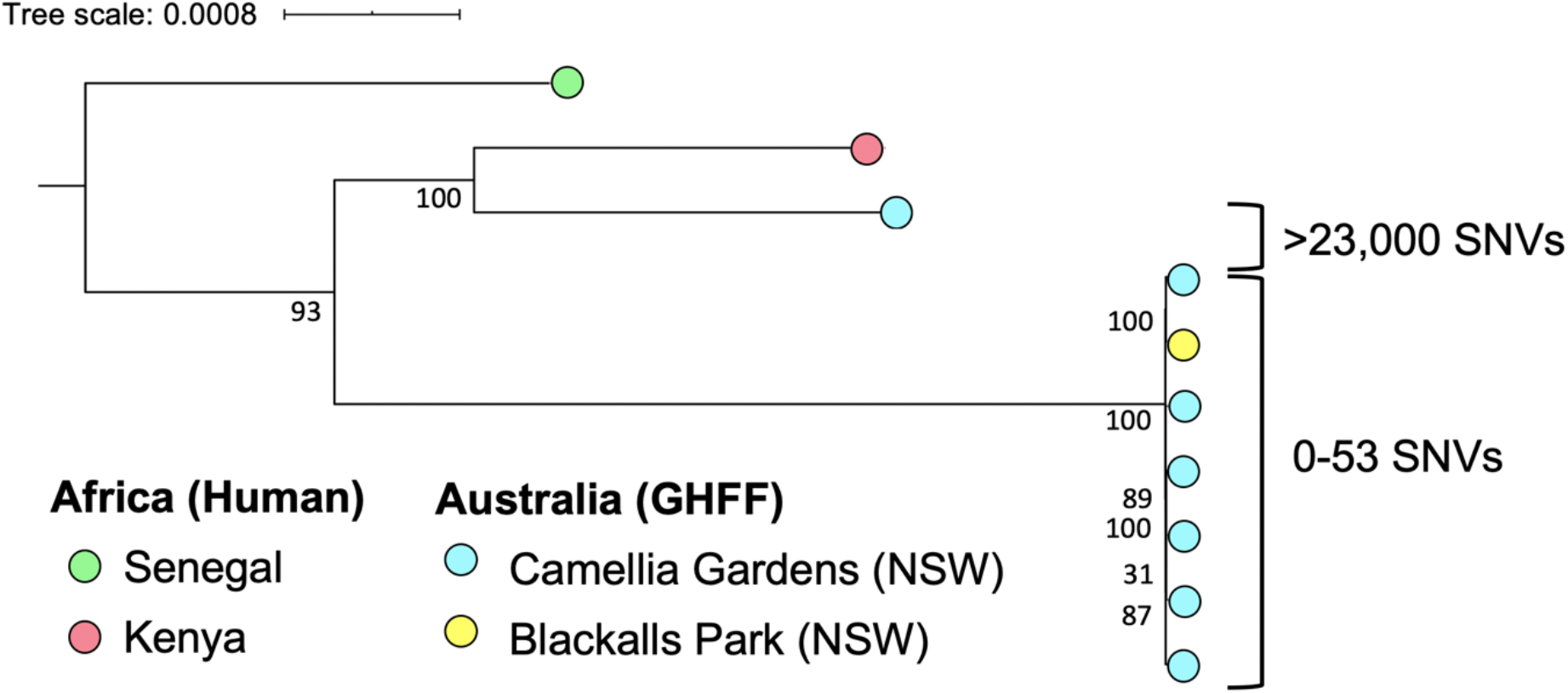
Core genome, outgroup-rooted (*K. quasivariicola* SRR5386043, not shown) maximum likelihood phylogenetic tree constructed using RAxML showing the relatedness of all GHFF and two human sourced *K. africana* isolates. SNVs were determined by read-mapping to the FF1003 chromosome (GenBank CP059391) using RedDog. Bootstrap values are based on 100 replicates. Scale bars indicate substitutions per site.

All *K. africana* isolates carried intrinsic *bla*_OKP-C-1_ genes which exhibited variations in their nucleotide sequences, however, the translated amino acid sequences were identical for all isolates. GHFF ST4938 *bla*_OKP-C-1_ sequences (*n*=7) had 2 SNVs compared to the ST3291 *bla*_OKP-C-1_ reference sequence in GenBank (MK161164). The single GHFF ST4939 isolate carried a *bla*_OKP-C-1_ nucleotide sequence with three SNVs compared to the reference sequence (MK161164). As noted above, all seven GHFF ST4938 isolates carried the yersiniabactin locus, which was absent in other *K. africana* lineages (Fig. 3).

## Discussion

The detection of two novel *K. africana* STs in GHFF effectively doubled the number of known lineages globally and increased the collective *K. africana* pool from two isolates to 10 isolates. This was a highly unexpected finding given that *K. africana* has previously only been isolated from humans in Africa [24]. The *K. africana* ST4938 bat isolates detected at two colonies geographically separated by 100km, exhibited low genetic diversity (0-53 SNVs). GHFF typically forage within 50 km from the colony location, however they can also travel longer distances between colonies over several days [15, 16]. These findings suggest a common source for the ST4938 by GHFF in the CG and BP colonies and possible horizontal microbial transmission between individuals roosting in close proximity [56]. The *K. africana* isolates were only detected in GHFF from a small geographical region and over a short temporal sampling period of four weeks, so it remains unclear if *K. africana* are part of the GHFF endemic microbiome. Diverse KpSC have been isolated from numerous environmental niches including plants, soil, insects and the aquatic environment [36]. Given the arboreal nature of wild GHFF, environmental reservoirs of *K. africana* such as fresh water or food resources, including native fruits and flowers, orchard fruits and insects, present possible sources [57, 58]. Wider sampling and high-resolution analysis of microbiome composition would assist in elucidating the frequency and ecology of *K. africana* in GHFF microbiomes.

In wild GHFF, the occurrence of KpSC isolates harbouring acquired ARGs was 1.2%, which was lower than the previously reported occurrence of class 1 integrons in wild GHFF (5.3%) [22]. Resistance to one critically important antimicrobial category, the fluoroquinolones, was detected in the GHFF *K. pneumoniae* isolates, whereas ESBL and carbapenemase activity was absent [3, 4]. These findings are comparable with an earlier study of AMR in Enterobacteriaceae from wild Australian mammals, which also found an absence of ESBL and carbapenemase producing *K. pneumoniae* and a low occurrence of resistance to other clinically important antimicrobials (range 0.0% to 10.1%) [59]. Recently, diverse strains of MDR *K. pneumoniae, Escherichia coli* and other Enterobacteriaceae harbouring plasmid-borne carbapenem resistance genes were detected in 40% of Australian silver gulls (*Chroicocephalus novaehollandiae*) at one site, demonstrating widespread transmission of ARGs in Australian wildlife [10]. In captive GHFF, no KpSC isolates harbouring acquired ARGs were found, which is in contrast to previous reports of a high occurrence of class 1 integrons in captive GHFF (41.2%) [22]. This absence of ARGs in KpSC isolates from captive GHFF group-housed in the same rehabilitation facility, may be explained by the widespread carriage (75% occurrence) of a single KpSC strain, *K. pneumoniae* ST5035, which did not harbour class 1 integrons or AMR genes.

Of the four *K. pneumoniae* STs shared by humans and wild GHFF, three have been reported as MDR *K. pneumoniae* in human clinical infections. ST1017 have been reported as MDR and ESBL producing human uropathogens [60] and isolated from human blood (Pasteur MLST, ID:1257), however, GHFF ST1017 isolates harboured only two acquired AMR genes (*dfrA14* and *qnrS1*) and were not considered MDR [61]. Two GHFF strains shared STs with carbapenemase producing strains reported to cause localised outbreaks ST105 [62] and ST661 [63], however neither GHFF isolate harboured any resistance genes. These findings suggest that although GHFF and humans share *K. pneumoniae* STs, the isolates detected in GHFF did not harbour MDR or virulence factors that are frequently associated with isolates causing MDR human nosocomial and community acquired infections.

Phylogenetic analysis suggests *K. africana* and *K. quasivariicola* are the closest relatives to the clinically important *K. pneumoniae* (*sensu stricto*) taxa [24]. The GHFF *K. africana* isolates exhibited characteristics associated with *K. pneumoniae* clinical infections, including the O2 serotype [55] and yersiniabactin [23]. Additionally, the human clinical *K. africana* isolate ST2831 carried diverse AMR genes, including the ESBL *bla*_CTX-M-15_ gene. These findings indicate *K. africana* strains are capable of acquiring virulence factors and AMR genes, and have the potential to be opportunistic pathogens of GHFF and humans [55].

The considerable variation of KpSC occurrence in GHFF from different locations may relate to differences in exposure to environmental sources of KpSC, such as water and food resources [36], and frequency of transmission between individuals, which is influenced by colony structure and density [56]. In addition to the apparent local clonal expansion of ST5035 in the captive Mylor GHFF, the clustering of three STs in the CG colony also suggests local clonal expansion, which may have contributed to the elevated occurrence of KpSC at CG compared to other wild colonies. Despite the detection of *K. pneumoniae* strain ST5035 in 75% of captive GHFF, it was notably absent in the parent colony (ABP), suggesting acquisition from a common source whilst in captivity, in combination with a high frequency of transmission between individuals. The captive GHFF were group-housed in an outdoor aviary with exposure to multiple potential sources of *K. pneumoniae* ST5035, including animals (poultry, dogs, captive kangaroos and other wildlife) and food (cultivated fruits) [36]. The stark contrast in the occurrence of *K. pneumoniae* between wild and captive GHFF, demonstrates widespread transmission of a potential bacterial pathogen in group-housed captive wildlife. Widespread acquisition (36.4% occurrence) of a class 1 integron harbouring the resistance gene *aadA2* has also been demonstrated in group-housed captive GHFF [22]. This potential for widespread dissemination of bacterial pathogens and AMR genes in group-housed captive GHFF should be considered as a potential health risk for both human carers and GHFF in rehabilitation programs.

Studies examining bacterial pathogens and AMR in wildlife typically focus on the potential for wildlife to act as bacterial reservoirs [64, 65] and largely ignore the potential impacts of anthropogenic pathogens colonising wildlife (reverse zoonosis) [66]. Each year, several thousand sick and injured GHFF require veterinary care [67], which may include antimicrobial therapy. The detection of fluoroquinolone resistant *K. pneumoniae* isolates capable of causing opportunistic infections in humans, and possibly in GHFF, is concerning, especially as enrofloxacin (a veterinary fluoroquinolone) is frequently prescribed to GHFF in care. Antimicrobial administration to GHFF carrying resistant isolates may result in a poor treatment response and prognosis for recovery, but also may increase dissemination to other GHFF in care, to human carers and into the environment post release. Additionally, increasing numbers of GHFF require veterinary care each year due to the cumulative impacts of wild fires, heat stress events, habitat loss and food shortages [67, 68].

This study has shown that GHFF carry diverse KpSC with varying frequency and can include organisms that are closely related to strains causing human disease, however, the data provided no evidence to suggest that GHFFs are a reservoir for the globally distributed MDR or hypervirulent *K. pneumoniae* lineages. We also report a new host and new geographic occurrence of *K. africana*, and four times the number of previously described *K. africana* isolates, including two new lineages, effectively doubling the number of known *K. africana* lineages. These findings highlight the necessity to further explore genome data for KpSC isolates from non-human niches, in order to expand our understanding of their ecology. Although the public health risks associated with KpSC isolates carried by GHFF are currently deemed low, our findings demonstrate the importance of surveillance of captive and wild GHFF for AMR and pathogenic bacteria, to meet the One Health strategy for managing AMR in Australia. This study further highlights the need for ecological studies of *K. pneumoniae* in diverse animal species to identify reservoirs of clinically important lineages.

## Acknowledgements

We wish to thank Terry Reardon, Ian Smith, Kathy Burbidge and the staff at the Animal Health Centre (Adelaide Zoo) for support in collecting samples from the Adelaide Botanic Park colony, Katrina Boardman and Fauna Rescue of South Australia for assistance in collecting samples from Mylor captive GHFF, and Juliane Schaer, Daniel Russell and Claudia Aguilar-Solis for assistance collecting samples from GHFF colonies in New South Wales. We thank the Institut Pasteur teams for the curation and maintenance of BIGSdb-Pasteur databases at http://bigsdb.pasteur.fr/. We thank Ryan Wick for assistance with the hybrid genome assembly of FF1003.

This work was supported by a Holsworth Wildlife Research Endowment (Equity Trustees Charitable Foundation and the Ecological Society of Australia) awarded to Fiona McDougall, a Lake Macquarie Council Environmental Research Grant awarded to Michelle Power and Fiona McDougall, and a University of Melbourne Early Career Researcher Grant awarded to Kelly Wyres.

