## Supplementary Tables S1 to S5 for "Novel strains of *Klebsiella africana* and *Klebsiella pneumoniae* in Australian Fruit Bats (*Pteropus poliocephalus*)"

**Table S1.** Primers used for *K. pneumoniae* MLST and class 1 integron screening. NA, Not applicable (standard PCR protocol).

| Target | Primer | Primer sequence 5’ to 3’ | Annealing temp (˚C) | Reference |
|---|---|---|---|---|
| *gapA* | gapA173 | TGAAATATGACTCCACTCACGG | 60 | [1] |
| | gapA181 | CTTCAGAAGCGGCTTTGATGGCTT | | |
| *mdh* | mdh130 | CCCAACTCGCTTCAGGTTCAG | 50 | [1] |
| | mdh867 | CCGTTTTTCCCCAGCAGCAG | | |
| *phoE* | phoE604.1 | ACCTACCGCAACACCGACTTCTTCGG | 50 | [1] |
| | phoE604.2 | TGATCAGAACTGGTAGGTGAT | | |
| *intI1* | HS463a | CTGGATTTCGATCACGGCACG | NA | [2] |
| | HS464 | ACATGCGTGTAAATCATCGTCG | | |
| *attI1* | HS458 | GTTTGATGTTATGGAGCAGCAACG | NA | [3] |
| *qacEΔ1* | HS459 | GCAAAAAGGCAGCAATTATGAGCC | | |

**Table S2.** Individual KpSC isolate whole genome sequence data accessioned in the NCBI Sequence Read Archive (SRA) under BioProject ID PRJNA646592.

| Isolate | ST | Species | BioProject | BioSample | read_accession |
|---|---|---|---|---|---|
| FF904 | 105 | *Klebsiella pneumoniae* | PRJNA646592 | SAMN15547544 | SRR12233577 |
| FF907 | 5032 | *Klebsiella variicola* ss. *variicola* | PRJNA646592 | SAMN15547547 | SRR12233610 |
| FF923 | 5034 | *Klebsiella pneumoniae* | PRJNA646592 | SAMN15547552 | SRR12233605 |
| FF930 | 4938 | *Klebsiella africana* | PRJNA646592 | SAMN15547533 | SRR12233580 |
| FF979 | 5033 | *Klebsiella pneumoniae* | PRJNA646592 | SAMN15547548 | SRR12233609 |
| FF983 | 5033 | *Klebsiella pneumoniae* | PRJNA646592 | SAMN15547549 | SRR12233608 |
| FF985 | 5033 | *Klebsiella pneumoniae* | PRJNA646592 | SAMN15547550 | SRR12233607 |
| FF986 | 5033 | *Klebsiella pneumoniae* | PRJNA646592 | SAMN15547551 | SRR12233606 |
| FF996 | 1017 | *Klebsiella pneumoniae* | PRJNA646592 | SAMN15547543 | SRR12233578 |
| FF1003 | 4938 | *Klebsiella africana* | PRJNA646592 | SAMN15547534 | SRR12233579 |
| FF1006 | 4938 | *Klebsiella africana* | PRJNA646592 | SAMN15547535 | SRR12233604 |
| FF1008 | 4938 | *Klebsiella africana* | PRJNA646592 | SAMN15547536 | SRR12233593 |
| FF1009 | 1017 | *Klebsiella pneumoniae* | PRJNA646592 | SAMN15547541 | SRR12233575 |
| FF1010 | 1412 | *Klebsiella pneumoniae* | PRJNA646592 | SAMN15547545 | SRR12233612 |
| FF1011 | 4938 | *Klebsiella africana* | PRJNA646592 | SAMN15547537 | SRR12233583 |
| FF1012A | 4919 | *Klebsiella pneumoniae* | PRJNA646592 | SAMN15547570 | SRR12233585 |
| FF1013 | 661 | *Klebsiella pneumoniae* | PRJNA646592 | SAMN15547568 | SRR12233587 |
| FF1014 | 4938 | *Klebsiella africana* | PRJNA646592 | SAMN15547538 | SRR12233582 |
| FF1016 | 1017 | *Klebsiella pneumoniae* | PRJNA646592 | SAMN15547542 | SRR12233574 |
| FF1017 | 4938 | *Klebsiella africana* | PRJNA646592 | SAMN15547539 | SRR12233581 |
| FF1019 | 4939 | *Klebsiella africana* | PRJNA646592 | SAMN15547540 | SRR12233576 |
| FF1023 | 5037 | *Klebsiella pneumoniae* | PRJNA646592 | SAMN15547571 | SRR12233584 |
| FF1043 | 4919 | *Klebsiella pneumoniae* | PRJNA646592 | SAMN15547569 | SRR12233586 |
| FF1076 | 5035 | *Klebsiella pneumoniae* | PRJNA646592 | SAMN15547553 | SRR12233603 |
| FF1077 | 5035 | *Klebsiella pneumoniae* | PRJNA646592 | SAMN15547554 | SRR12233602 |
| FF1078 | 5035 | *Klebsiella pneumoniae* | PRJNA646592 | SAMN15547555 | SRR12233601 |

| | | | | | |
|---|---|---|---|---|---|
| FF1079 | 5035 | *Klebsiella pneumoniae* | PRJNA646592 | SAMN15547556 | SRR12233600 |
| FF1080 | 5035 | *Klebsiella pneumoniae* | PRJNA646592 | SAMN15547557 | SRR12233599 |
| FF1081 | 5035 | *Klebsiella pneumoniae* | PRJNA646592 | SAMN15547558 | SRR12233598 |
| FF1082 | 5035 | *Klebsiella pneumoniae* | PRJNA646592 | SAMN15547559 | SRR12233597 |
| FF1083 | 5036 | *Klebsiella pneumoniae* | PRJNA646592 | SAMN15547546 | SRR12233611 |
| FF1084 | 5036 | *Klebsiella pneumoniae* | PRJNA646592 | SAMN15547560 | SRR12233596 |
| FF1085 | 5035 | *Klebsiella pneumoniae* | PRJNA646592 | SAMN15547561 | SRR12233595 |
| FF1086 | 5035 | *Klebsiella pneumoniae* | PRJNA646592 | SAMN15547562 | SRR12233594 |
| FF1088 | 5035 | *Klebsiella pneumoniae* | PRJNA646592 | SAMN15547563 | SRR12233592 |
| FF1089 | 5035 | *Klebsiella pneumoniae* | PRJNA646592 | SAMN15547564 | SRR12233591 |
| FF1091 | 5035 | *Klebsiella pneumoniae* | PRJNA646592 | SAMN15547565 | SRR12233590 |
| FF1092 | 5035 | *Klebsiella pneumoniae* | PRJNA646592 | SAMN15547566 | SRR12233589 |
| FF1094 | 5035 | *Klebsiella pneumoniae* | PRJNA646592 | SAMN15547567 | SRR12233588 |

**Table S3.** Complete genome assemblies for isolate FF1003 accessioned in GenBank.

| Isolate | Species | BioProject | BioSample | Localid | Accession |
|---|---|---|---|---|---|
| FF1003 | *Klebsiella africana* | PRJNA646592 | SAMN15547534 | chromosome | CP059391 |
| FF1003 | *Klebsiella africana* | PRJNA646592 | SAMN15547534 | plasmid_1 | CP059392 |
| FF1003 | *Klebsiella africana* | PRJNA646592 | SAMN15547534 | plasmid_2 | CP059393 |

**Table S4.** Antibiotic disks used for EUCAST susceptibility testing.

| Antibiotic and disk content (µg) | OXOID disk | Antibiotic category and generation | Breakpoint |
|---|---|---|---|
| Ampicillin (10 µg) | AMP10 | ß-lactam/Penicillins | EUCAST |
| Amoxicillin/Clavulanic Acid (30 µg) | AMC30 | ß-lactam/Penicillins + ß-lactamase inhibitors | EUCAST |
| Cephalexin (30 µg) | CL30 | ß-lactam/Cephalosporins/1st Generation (Non-extended spectrum cephalosporin) | EUCAST |
| Cefotaxime (5 µg) | CTX5 | ß-lactam/Cephalosporins/3rd Generation (Extended-spectrum cephalosporin) | EUCAST |
| Imipenem (10 µg) | IPM10 | ß-lactam/Carbapenems | EUCAST |
| Nalidixic acid (30 µg) | NA30 | Quinolones-Fluoroquinolones/ 1st Generation Quinolone | CLSI |
| Ciprofloxacin (5 µg) | CIP5 | Quinolones-Fluoroquinolones/ 2nd Generation Fluoroquinolone | EUCAST |
| Trimethoprim (5 µg) | W5 | Trimethoprim (Folate pathway inhibitors) | EUCAST |
| Trimethoprim/Sulfamethoxazole (30 µg) | SXT30 | Trimethoprim + Sulfonamide (Folate pathway inhibitors) | EUCAST |
| Amikacin (30 µg) | AK30 | Aminoglycosides | EUCAST |

**Table S5.** Phenotypic antimicrobial susceptibility testing results for all GHFF KpSC isolate strains (*n*=13). Amoxicillin resistance was determined as growth in LB broth supplemented with 10 mg/L amoxicillin (EUCAST MIC breakpoint for resistance >8 mg/L. All other antibiotic susceptibility testing was performed using the EUCAST disk diffusion method. IBLRG; Intrinsic beta-lactamase resistance genes. R; Resistant. I; Intermediate resistance. S; Susceptible. AMX; Amoxicillin. AMP; Ampicillin. AMC; Amoxicillin plus clavulanic acid. CL; Cephalexin. CTX; Cefotaxime. IPM; Imipenem. NA; Nalidixic acid. CIP; Ciprofloxacin. W; Trimethoprim. SXT; Trimethoprim/Sulfamethoxazole. AK; Amikacin.

| IBLRG | Other ARGs | ST | No. isolates | AMX | AMP | AMC | CL | CTX | IPM | NA | CIP | W | SXT | AK |
|---|---|---|---|---|---|---|---|---|---|---|---|---|---|---|
| SHV-1^ | - | 105 | 1 | R | R | S | S | S | S | S | - | S | S | S |
| SHV-1 | - | 1412 | 1 | R | R | S | S | S | S | S | - | S | S | S |
| SHV-1^ | - | 5037 | 1 | R | R | S | S | S | S | S | - | S | S | S |
| SHV-1^ | - | 5036 | 1 | R | R | S | S | S | S | S | - | S | S | S |
| SHV-1 | - | 5033 | 4 | R | R | S | S | S | S | S | - | S | S | S |
| SHV-11.v1^ | *dfrA14, qnrS1* | 1017 | 3 | R | R | S | S | S | S | I | R | R | S | S |
| SHV-27 | - | 661 | 1 | R | R | S | S | S | S | S | - | S | S | S |
| SHV-27^ | - | 4919 | 2 | R | R | S | S | S | S | S | S | S | S | S |
| SHV-110^ | - | 5035 | 15 | R | R | S | S | S | S | S | - | S | S | S |
| SHV-110* | - | 5034 | 1 | R | R | S | S | S | S | S | - | S | S | S |
| LEN-2^ | - | 5032 | 1 | R | R | S | S | S | S | S | - | S | S | S |
| OKP-C-1^ | - | 4938 | 7 | R | R | S | S | S | S | S | - | S | S | S |
| OKP-C-1^ | - | 4939 | 1 | R | R | S | S | S | S | S | - | S | S | S |

^ Identical translated amino acid sequence (nucleotide sequence variant)

* Nucleotide sequence variant (closest match)
